# Small molecule chaperones facilitate the folding of long non-coding RNA G

**DOI:** 10.1101/2023.04.04.535601

**Authors:** Pauline Lejault, Louis Prudent, Michel-Pierre Terrier, Jean-Pierre Perreault

**Affiliations:** Department of biochemistry and functional genomics, Pavillon de recherche appliquée sur le cancer, Université de Sherbrooke, Sherbrooke, Québec, Canada, J1E 4K8

## Abstract

RNA G-quadruplexes (rG4) have recently emerged as major regulatory elements in both mRNA and non-coding RNA. To investigate the biological roles of the rG4 structures, chemists have developed a variety of highly specific and potent ligands. All these ligands bind to the rG4 by staking on their top, and the specificity of binding is demonstrated in comparison to other structures such as duplex or three-way junctions. It remains unclear whether rG4-ligands merely stabilize fully formed rG4 structures, or if they actively participate in the folding of the rG4 structure through their association with an unfolded RNA sequence. In order to access the innate steps of ligand-rG4 association and mechanisms, robust *in vitro* techniques, including FRET, electrophoretic mobility shift assay and reverse transcriptase stalling assays, were used to examine the capacity of five well-known G4 ligands to induce rG4 structures derived from either long non-coding RNAs of from synthetic RNAs. It was found that both PhenDC3 and PDS induce rG4 formation in unfolded single RNA strands. This discovery has important implications for the interpretation of RNA-seq experiments. Overall, *in vitro* data that can assist biochemists in selecting the optimal G4-ligands for their RNA cellular experiments are presented, while also considering the effects induced by these ligands of the rG4.

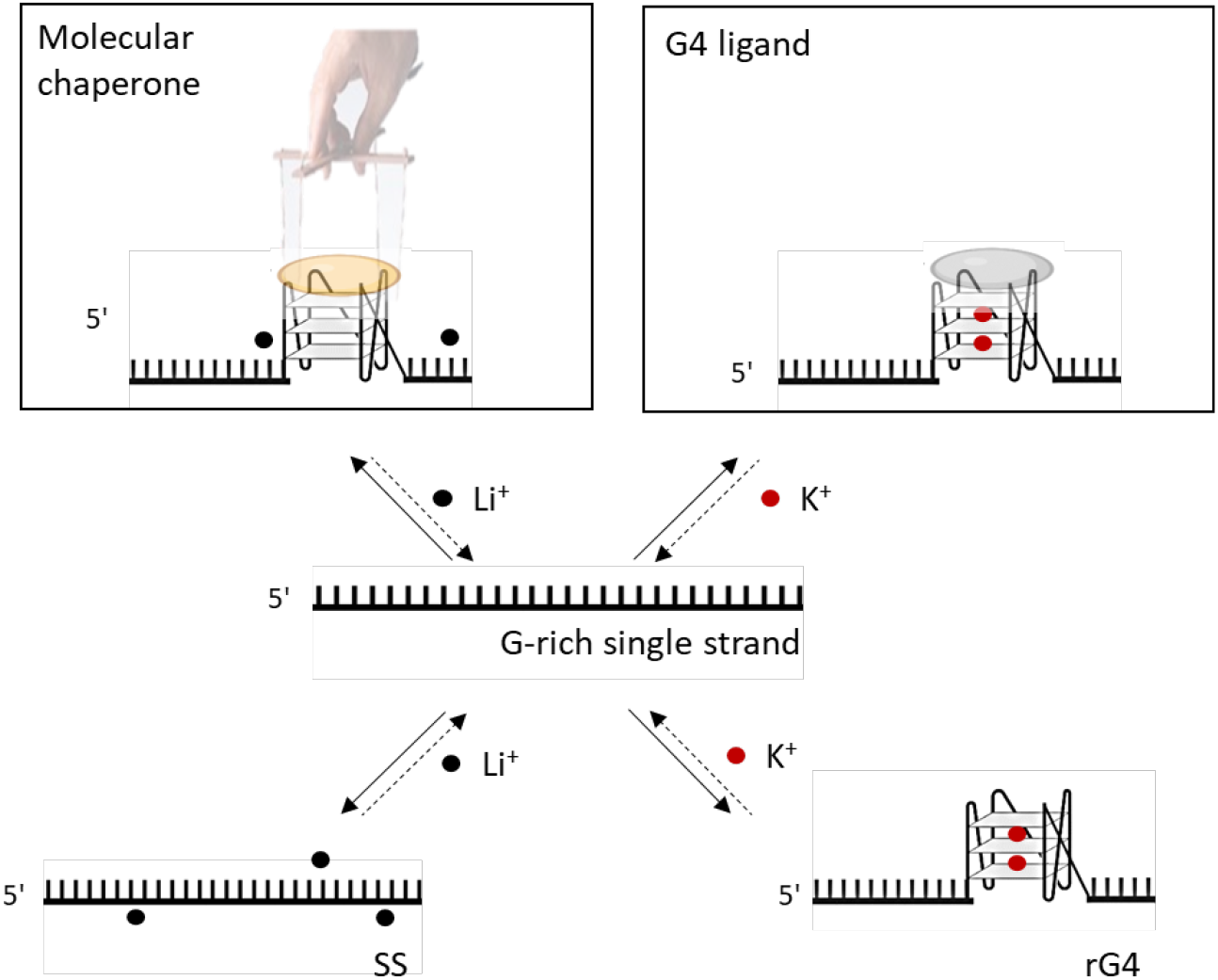

**Graphical abstract:** A schematic representation of the potential structures that may arise from unfolded RNA that is rich in G nucleotides. rG4 can be stabilized by K^+^ either with or without G4 ligand and can be induced by a molecular chaperone in the presence of Li^+^.

## Introduction

Through their affinity and selectivity for G-quadruplexes (G4s), G4-ligands are invaluable tools with which to investigate the plethora of G4 biological functions in cells^1,2^. In a typical biochemical strategy, G4 ligands are used to reversibly disrupt the processes that G4s are involved in, thus providing readouts amenable to mechanistic interpretations. G4 ligands have been instrumental in uncovering the regulatory roles of G4s in both gene expression (at both the transcriptional (dG4) and the translational levels (rG4)) and in key cellular processes (e.g. telomeric equilibrium, replication origin, autophagy, polyadenylation, splicing, etc.)^3–6^. This also explains why G4-involving cellular events are now being actively investigated as targets for chemotherapeutic interventions, according to an innovative strategy relying on the use of nucleic acid structure-specific binders^7,8^.

Amongst the thousands of G4 ligands, the most popular are undoubtedly PhenDC3 and pyridostatin (PDS) (Figure 1A)^9^. These two pyridodicarboxamide (PDC) derivatives have been widely used by many research teams worldwide as they have shown remarkable potential in binding DNA G4s. For instance, PhenDC3, which was developed by Cian *et al*. ^10^, was used at both the genomic and the transcriptomic levels to block telomerase progression and repress the translation of an oncogenic mRNA (TRF2)^11–13^, respectively. On the other hand, pyridostatin (PDS), which was developed by Rodriguez *et al*. ^14^, was used to identify cellular pathways involving G4s in the entire genome. PDS was also found to be an invaluable *in vitro* molecular tool for sequencing-based methods such as G4-seq^15^ and rG4-seq^16^. The structural simplicity of both PhenDC3 and PDS makes them amenable to chemical derivatization that endows them with new functionalities. For instance, PDS was recently linked to a fluorophore (SiR-PyPDS ^17^) so as to be able to follow with the aim to isolate bound G4s *via* affinity precipitation protocols^18^. Both PhenDC3 and PDS were conjugated to alkyne appendages so that the resulting derivatives (PhenDC3-alk^19^ and PDS-*α* ^20^) could be used for bioorthogonal cellular manipulations (notably *in situ* click imaging)^21^.

**Figure 1:**
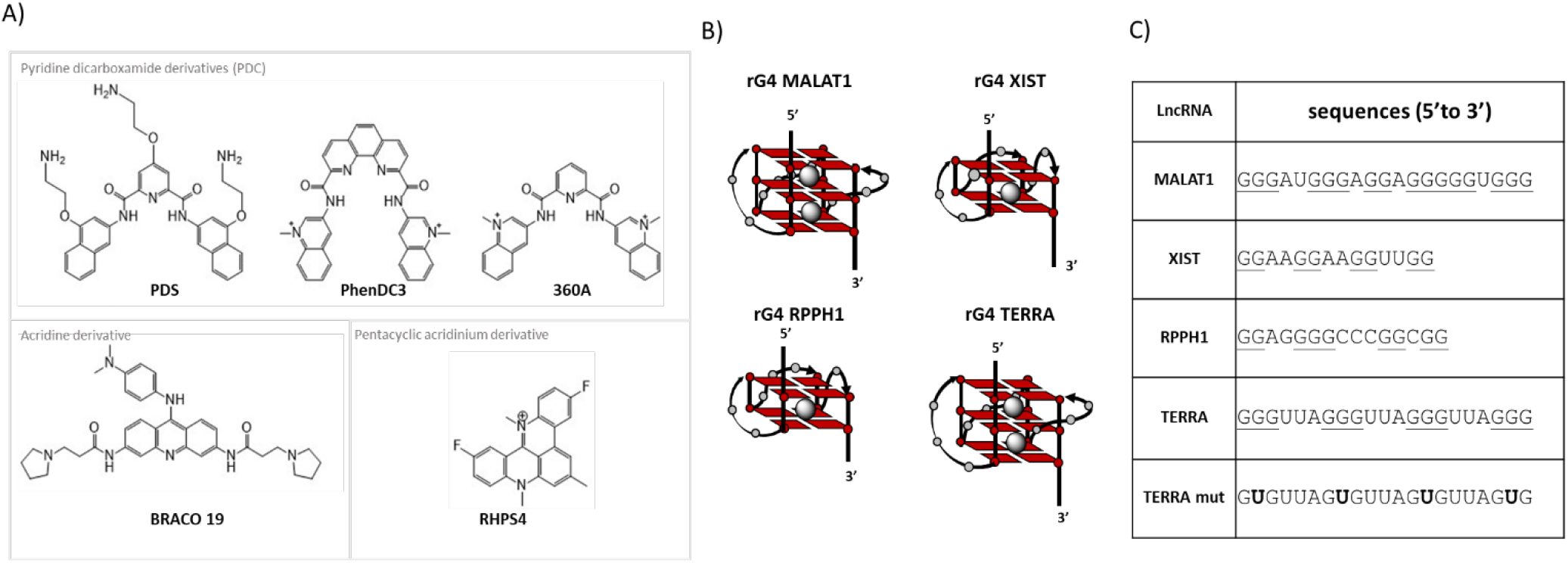
Ligands and rG4 derived from lncRNA-derived used in this study. A) Chemical structures of the G4 ligand evaluated as potential molecular chaperones. B) A schematic representation of potential structures of G4 structure folded by MALAT1, XIST, RPPH1 and TERRA. C) The minimal sequences that were used by BioTASQ to fold the lncRNA into an rG4 trap.

Interestingly, the very first prototypes of PDC were 307A^22^ and 360A^23^ (Figure 1A). Initially identified by Sanofi-Aventis, these compounds turned out to be efficient tools with which to demonstrate the existence of G4s in telomeres (*via* the use of tritiated 360A^23^ which influences the telomeres’ integrity^24,25^). The chemistry around the 360A motif is also well-developed. For example, this ligand was conjugated to biotin (PDC-biotin) for the SELEX-type selection of G4 motifs^26^, to a fluorophore (PDC-TO) in order to permit visualization of G4 formation *in vitro*^27^ and to cross-linking moieties (benzophenone and 4-azido-benzoic acid, PDC-XL) to identify cellular G4 partners^28^.

Less studied in the last decade, Braco-19 and RHPS4, respectively acridine and petacycle acridinium derivatives, are also known as potent telomerase inhibitors^29^. Braco-19 has been widely applied to the targeting of G4s from viral genomes^8^. Elsewhere, Yang et al have shown notable differences in the modulation of non-coding RNAs’ profiles following the use of these G4 ligands by the G4RP-seq method^30^.

These transcriptomic landscape studies embolden the future use of G4-stabilizing agents to help in understanding rG4 biology. It is therefore intriguing that the exact mechanism by which these G4 ligands interact with, or induce G4s, in RNA is not yet fully deciphered. Inspired by pioneering work done in the dG4 context by both Mergny *et al*.^31^ and Birkedal *et al*.,^32^, it is demonstrated here that both PhenDC3 and PDS can act as molecular chaperones and promote the formation of rG4 *in vitro;* while RHPS4 and Braco19 interact mostly with folded G4s, which has important consequences in terms of G4 ligand selection for cellular experiments.

## Materials and methods

All oligonucleotides used in this study were purchased from Integrated DNA technology (IDT) and were stored at -20°C as 100 to 1000 µM stock solutions in deionized water. The actual concentration of these stock solutions was determined by spectrophotometry at 260 nm. PhenDC3 and Pyridostatin (PDS) were purchased from Sigma; Tmpyp4, Braco 19 and RHPS4 were purchased from Bio-techne; and, 360A was synthesized according to the literature^33^. All G4 ligands were dissolved in DMSO at a concentration of 2.5 mM prior to storage. These stock solutions were then diluted with deionized water to the appropriate concentration for the experiment in question. MmuLV reverse transcriptase was produced and purified by the protein purification platform of the Université de Sherbrooke.

### *In vitro* transcription

DNA templates containing the T7 RNA promoter were designed (see Figure S1). Double-stranded DNAs were produced *via* a PCR filling strategy using the T7 promoter oligonucleotide. Then, an *in vitro* transcription was performed as described previously^34^. Subsequently, the proteins were removed by phenol/chloroform extraction followed by ethanol precipitation of the RNA. The transcripts were then purified using denaturing (8 M urea) 5% to 10% polyacrylamide gels (19:1), depending on the length of the transcripts, followed excision of the gel fragment containing the transcript. The transcripts were then eluted from the gel slices in buffer TEL800 (10 mM Tris–HCl pH 7.5, 1 mM EDTA, 800 mM LiCl) and ethanol precipitated^35^. Lastly, the resulting RNA pellets were washed with 70% ethanol, air-dried, and dissolved in RNase-free water. RNA concentrations were determined by spectrometry at 260 nm.

### FRET melting experiments

FRET experiments were performed in 96-well plates using an Mx3005P qPCR machine (Agilent) equipped with FAM filters (λ_ex_ = 492 nm; λ_em_ = 516 nm). Each experiment was performed in a final volume of 100 μL of 10 mM cacodylate Li buffer with 0.2 μM of double-labeled (Fam/Tamra) RNAs (Eurogentec) and either with or without 5 molar equivalents (mol.eq) of the G4 ligands (*i*.*e*. PhenDC3, PDS, Braco19, RHPS4, and 360A). After the first equilibration step (25°C, 30 s) of the FRET melting experiments, a stepwise increase of 1°C every 30 s for 65 cycles was performed to reach 90°C.Measurements were performed after each cycle. The emission of FAM was normalized (0 to 1), and the T_m_ was defined as the melting temperature for which the normalized emission was 0.5. ΔT_1/2_values were calculated as follows: ΔT_1/2_ = [T_1/2_(RNA+ligand)-(T_m_(RNA alone)] and are the mean values of three experiments. Final data were analyzed using both Excel (Microsoft Corp.) and Prism 9-Graph Pad. The melting curves are presented in Figure S2.

### Reverse transcriptase stalling assays

The reverse transcriptase stalling (RTS) assays were performed using *in vitro* synthesized RNA transcripts (see Figure 2 for the RNA structures and Figure S1 for the oligonucleotides) as described previously with the modifications described below^35^. Each experiment was performed in the presence of 4.4 pmol of RNA and 5 pmol of Cy5-labeled primer in RTS buffer (50 mM Tris, 5 mM MgCl_2,_ and 75 mM of either LiCl or KCl) in a final volume of 7.5 μL. A pairing step of 3 min at 75°C containing the RNA, the labeled primer and the RTS buffer without MgCl_2_ was performed. The temperature of the mixture was then decreased to 37 °C for 5 min and 0.5 μL of 0.1M DTT, 0.5 μL of 10 mM dNTPs and 1 μL of 25 mM MgCl_2_ were then added. Then, 1 μL of either 44 μM ligand (10 equiv/RNA) or water was added. To generate the ladders, 1 μL of 10 mM ddNTP (di-deoxyribonucleotide) was also incorporated into the reaction at this moment. Finally, 0.5 μL of purified MMuLV reverse transcriptase was then added to each tube and the reactions were incubated for 15 min at 45°C. The enzyme was then inactivated by adding 0.75 μL of 2N NaOH per sample and heating at 90°C for 10 min. Fifteen microliters of loading denaturing buffer (98% formamide, 10 mM EDTA) were added to each sample and they were then either stored at 4°C or were used for the denaturing gel migration. Ten microliters were loaded onto denaturing (8M Urea) 12% polyacrylamide gels (19:1) which were electrophoresed for 2 h at 50 W. The final images were generated using a GE Typhoon FLA 9000 fluorescence scanner (λ_ex_=635 nm) and are presented in Supplementary Figures S5-S7). The “stop” bands were quantified using the ImageJ software by dividing the intensity of the stop region by the combined intensities of the stop region and stop corresponding to the full length. The relative ligand effect was obtained by dividing the intensity of the stop percentage observed in the presence of the ligand (i.e., either the Li^+^ or the K^+^ condition) with that observed in the presence of the RNA, but without any ligand being present (i.e. either the Li^+^ or the K^+^ condition). See Figure S4 for further details. All experiments were performed at least in duplicate.

**Figure 2:**
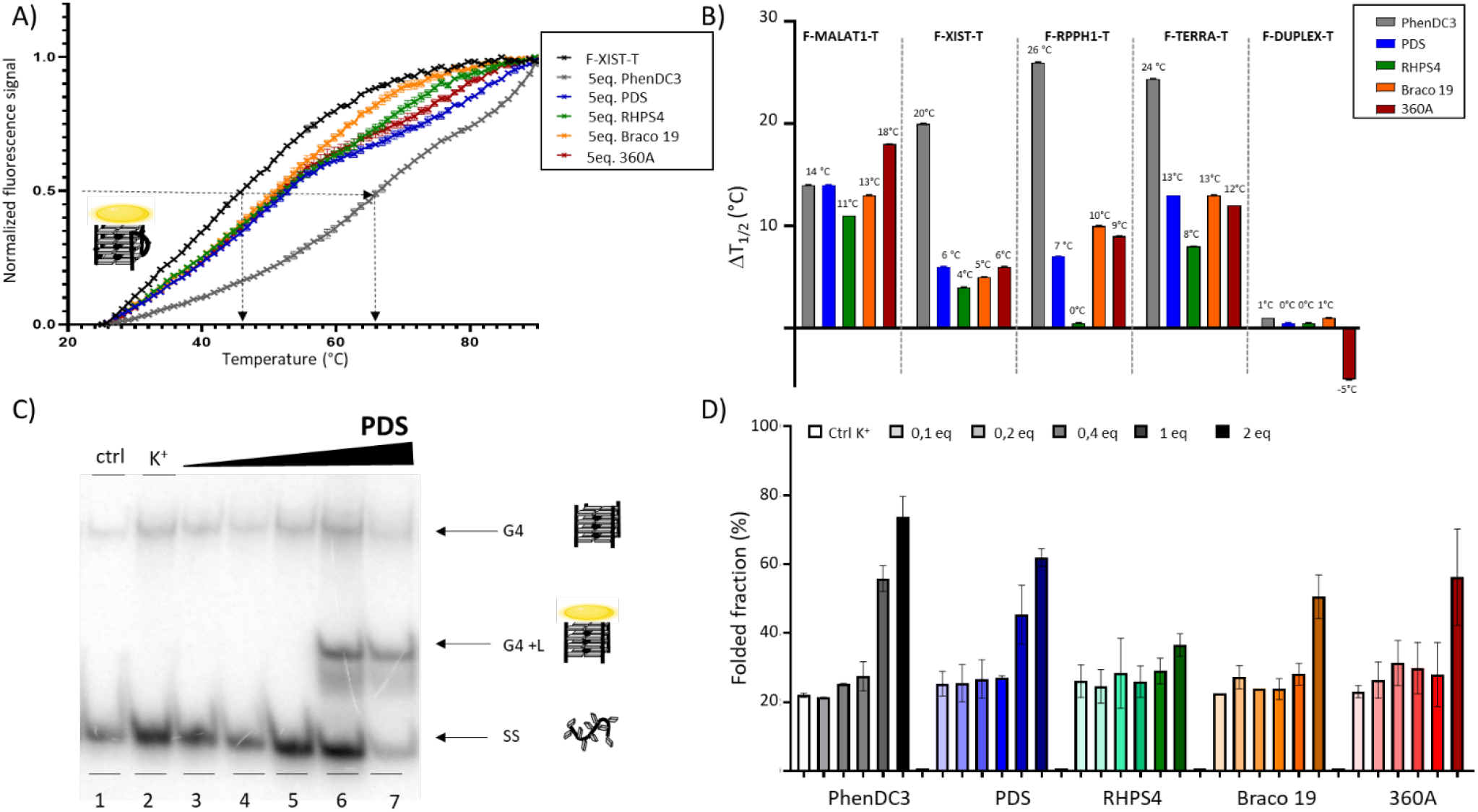
Examining the binding affinity and selectivity of various ligands on the rG4s. A) Example of the FRET-denaturing spectra obtained with F-xist-T either alone (black line), or in the presence of 5 mol. eq of G4 ligands in Li^+^ buffer. B) ΔT_1/2_ induced by 1 µM of PhenDC3, PDS, RHPS4, Braco 19 or 360A. C) Example of a native gel shift assay performed with ^32^P labelled-A_10_G_4_A_10_ (20uM) in the presence of increasing amounts of PDS (0,1;0,2; 0,4; 1 and 2 mol. eq. D) Compilation of the EMSA folded fractions observed with increasing amounts of the five ligands.

### NMM fluorescence assays

Fluorescence assays were performed as described previously ^36^. Either the G4 folding RNAs, or the control with a G/A mutant (200 pmol), were added to the folding buffer (20 mM Li-cacodylate pH 7.5 and 100 mM of either LiCl or KCl). The reactions were then heated at 70°C for 5 min before being slowly cooled to room temperature over a period of 1 h. The appropriate buffer (20 mM Li-cacodylate pH 7.5, 20 mM MgCl_2_, and 100 mM of either LiCl or KCl) was then added to a final volume of 100 μL. Next 2.5 eq/RNA of N-Methyl-Mesoporphyrin IX (NMM) (Frontier Scientific Inc., Logan, Utah) was added and the reaction incubated in the dark for 5 min at room temperature in a 10 mm quartz cuvette. The fluorescence intensity was then monitored using a Hitachi F-2500 fluorescence spectrophotometer with an excitation wavelength of 399 nm, and the emission spectra were recorded between 500 and 650 nm. The fluorescence at 605 nm was used for quantification (Figure S8). All NMM assays were performed at least in duplicate.

### Radiolabeling and electrophoretic mobility shift assays

RNAs were labeled with γ^32^P-ATP using T4 polynucleotide kinase (PNK) according to the manufacturer conditions (New England Biolabs). Radioactive RNAs were separated from any non-incorporated γ^32^P-ATP via denaturing 15% polyacrylamide (19:1) gel electrophoresis. The RNAs were eluted from the gel and ethanol precipitated as described above in the “*In vitro* Transcription” section. As described previously^31^, all samples were prepared in a final volume of 20 μL containing 20 μM of cold RNA, a negligible amount of radiolabeled RNA and increasing amounts of the G4-ligands (0,1 to 40 μM). Ten millimolar lithium cacodylate buffer (pH 7.2) without K^+^ was used for the control reaction, while the same buffer plus 10 mM KCl was used for all other conditions. Each sample was then heated for 5 min at 80**°**C and then slowly cooled at room temperature over a period of 2 h. Finally, 2 µL of loading buffer (50% sucrose and 50% H_2_0 without tracking dye) and 4 µL of each sample were mixed and then loaded on non-denaturing 20% polyacrylamide-bisacrylamide (29:1) gels. The electrophoretic mobility shift assays (EMSA) were performed in 1×TBE (Tris-borate EDTA buffer) containing 10 mM KCl (pH 8.3) for 15 min at 7 W followed by 2 h at 15 W, all at 4°C. After electrophoresis, the gels were dried at 50**°**C for 50 min and exposed to a phosphor screen that was subsequently scanned with a Typhoon™ trio imaging system (GE Healthcare). The results are presented in Figure S3. EMSA were performed and quantified at least in duplicate using the ImageJ software.

## Results and discussions

Several studies have demonstrated, through RNA sequencing, the formation of rG4 structures in the transcriptome and have reported the use of small molecular tools with which to study the dynamics and visualization of these structures in living cells. Although most G4 ligands have been tested as stabilizers of G4 structures, meaning that they provoke the thermal stabilization of these structures, the impact of G4 ligands on rG4 folding is still not fully understood. Consequently, it decided to evaluate a panel of five representative G4 ligands (PhenDC3, PDS, Braco19, RHPS4 and 360A; see Figure 1A) using several biophysical techniques. The aim was to determine whether G4 ligands can act as molecular chaperones by not only binding to rG4s, but also by inducing their formation. The investigation initially focused on four rG4 structures derived from well-known long non-coding RNAs (lncRNA): MALAT1, XIST, RPPH1 and TERRA, all of which have previously been reported as *in vitro* G4-forming sequences^37–40^ (Figure 1B-C). Interestingly, when compared to untreated cells, the first three have been found to be enriched in the global transcriptomic profile of MCF7 cells treated with G4 ligands (Braco19 and RHPS4) by the selective G4 molecular tool BioTASQ ^30^. The telomeric transcript TERRA, which is one of the most studied lncRNAs required for telomere integrity^41^, was also included in this study. The structures of each potential rG4 identified are depicted in Figure 2B. The investigation was centered around the G-rich segment of the lncRNA molecules, specifically the smallest portion capable of forming an rG4 structure (Figure 2C). Based on their sequences, RPPH1 and XIST are presented as non-canonical rG4s since they were found to form G4s consisting of only 2 quartets. On the other hand, TERRA and MALAT1 were classified as canonic rG4s due to their formation of 3-quartet rG4 structures.

### Investigating the folding and assembly properties of five well-studied G4 ligands

Historically, FRET melting assays have been performed in G4-favorable buffers containing KCl or NaCl with a stepwise increase in temperature from 20°C to 90°C to determine the thermal stability gain due to the addition of G4 ligands on either rG4s or dG4s (doubly labeled) ^42^. Based on this, the classic FRET melting assays were performed in unfavorable salt conditions (e.g., 1 mM cacodylate and in the presence of 10 mM Li^+^) for rG4 formation. As illustrated in Figure 2A, the apparent affinity of the G4-ligands was evaluated at 5 mol. eq of RNA. The stabilization was found to be quite high for PhenDC3 (ΔT_1/2_ up to 14°C) and to be selective (ΔT_1/2_ = 1°C with f-duplex-t) (Figure 2B). PDS, Braco 19 and 360A have an apparent affinity that is comparable. Conversely, RHPS4 does not display good affinity and has a lower melting temperature (ΔT_1/2_ lower than 13°C). These results show that all these candidates interact with unfolded G4s, but that PhenDC3 is the best stabilizing G4-ligand for most of the rG4 structures tested.

The reverse thermal curve, a renaturation caused by decreasing the temperature from 90°C to 20°C was also examined to evaluate the ability of the G4 ligands to induce intramolecular G4 formation upon binding to an unfolded G4-forming sequence (Figure S2). The results confirmed the good folding profile of PhenDC3, but the ΔTr was surprisingly negative with F-Malat1-T, possibly due to the polymorphism of f-Malat1-t. Despite the importance of rG4s, the folding mechanisms remain hard to investigate *in vitro*, since the rG4s are more stable and dynamic than their dG4 counterparts^43^. The folding and unfolding states of intramolecular rG4s without G4 ligands are multiple and complex (e.g. they are capable of forming hairpin, triplex and double hairpin states), which may explain the diversity of transitions obtained by FRET-renaturing assays (Figure S2)^44^.

The simple tetramolecular rG4 models offer a good opportunity with which to evaluate the assembling ability of the G4 ligands. This has been previously done for DNA G4 using 360A ligand^45^. Although limited, the formation of tetramolecular rG4s can be particularly relevant as G-rich sequences of tRNAs have been shown to assemble into intermolecular rG4s, leading to the formation of stress granules^46^. To do so, the formation of rG4s have been directly determined by EMSA using radiolabeled single-stranded RNA molecules (A_10_G_5_A_10_) under native conditions (i.e., in a non-denaturing K^+^ buffer). The salt strength (10 mM K^+^) and RNA concentration (20 μM) have been diminished as compared to the earlier protocol in order to illustrate the impact of the G4 ligands on the G4 association process^47^. After a slow cooldown in the presence of K^+^, a partial rG4 formation is observed (around 21%), and the single-stranded RNA remains the main population (Figure 2C, lane 2). Increasing concentrations of ligands had diverse impacts depending on their chemical structure. Unexpectedly, the ligand/rG4 complex formed with PhenDC3, PDS and Braco 19 migrated faster than the rG4 alone. This might be due to the molecular weight/charge and volume of the complex (see Figure 2C and Figure S3). In accordance with the FRET-renaturing experiments, both RHPS4 and Braco 19 induce the formation of rG4 to a lesser extent than do the other compounds. The relative folded fraction (including the rG4 formation and the complex ligand/rG4) does not exceed 36% and 50% with 2 mol. eq of RHPS4 and Braco 19, respectively (see Figure 2D). In comparison, both PhenDC3 and PDS lead to the majority of either the rG4 or the rG4 complex being folded, with 73% and 62% of the folded fractions being observed for the same molecular ratio conditions, respectively. Altogether, the data collected through these complementary techniques provided interesting insights into the G4-chaperone properties of the G4 ligands, but is insufficient to select the most versatile tools to either help the formation or to only stabilize existing rG4s.

### G4 induction by PhenDC3 and PDS impedes reverse transcriptase processivity

The RNA G4-mediated reverse transcriptase stalling (RTS) assays developed by *Kwok et al* ^35^, which were originally designed to identify rG4s and map their structures at a nucleotide resolution as demonstrated by the occurrence of an RT-stall in the presence of K^+^, were adapted. More precisely, this approach was used to identify the position of the rG4 induced by the ligand in the presence of Li^+^, as well as to evaluate the strength of the folded structure in the absence of K^+37^. To achieve this, a double-structured RNA which possessed a 5’-hairpin that was used as an internal RTS control, a 3’-hairpin for primer binding and the guanine-rich sequence of interest located in the middle was used. When the Cy5-DNA primer is hybridized to the RNA template, the reverse transcriptase can mediate the synthesis of a DNA molecule along the G-rich RNA template. In the presence of an excess of K^+^ (150 mM), the assay showed specific stalls at rG4 sites, causing an accumulation of truncated cDNAs (see Figure 3A). The RTS assays were performed in the presence of either Li^+^ or K^+^, and either with or without the G4 ligands to confirm G4 formation in the double hairpin context of the four rG4 sequences. The stall was observed at the predicted first track of guanines, with the intensities varying depending on both the sequence and the G4 ligand used (Figures S5-7). First, stop quantifications were meticulously performed in order to compare the G4 formation of the candidates under both favorable (K^+^) and unfavorable (Li^+^) salt conditions in the absence of any ligands (Figure 3B). The quantification method used is described in Figure S4.

**Figure 3:**
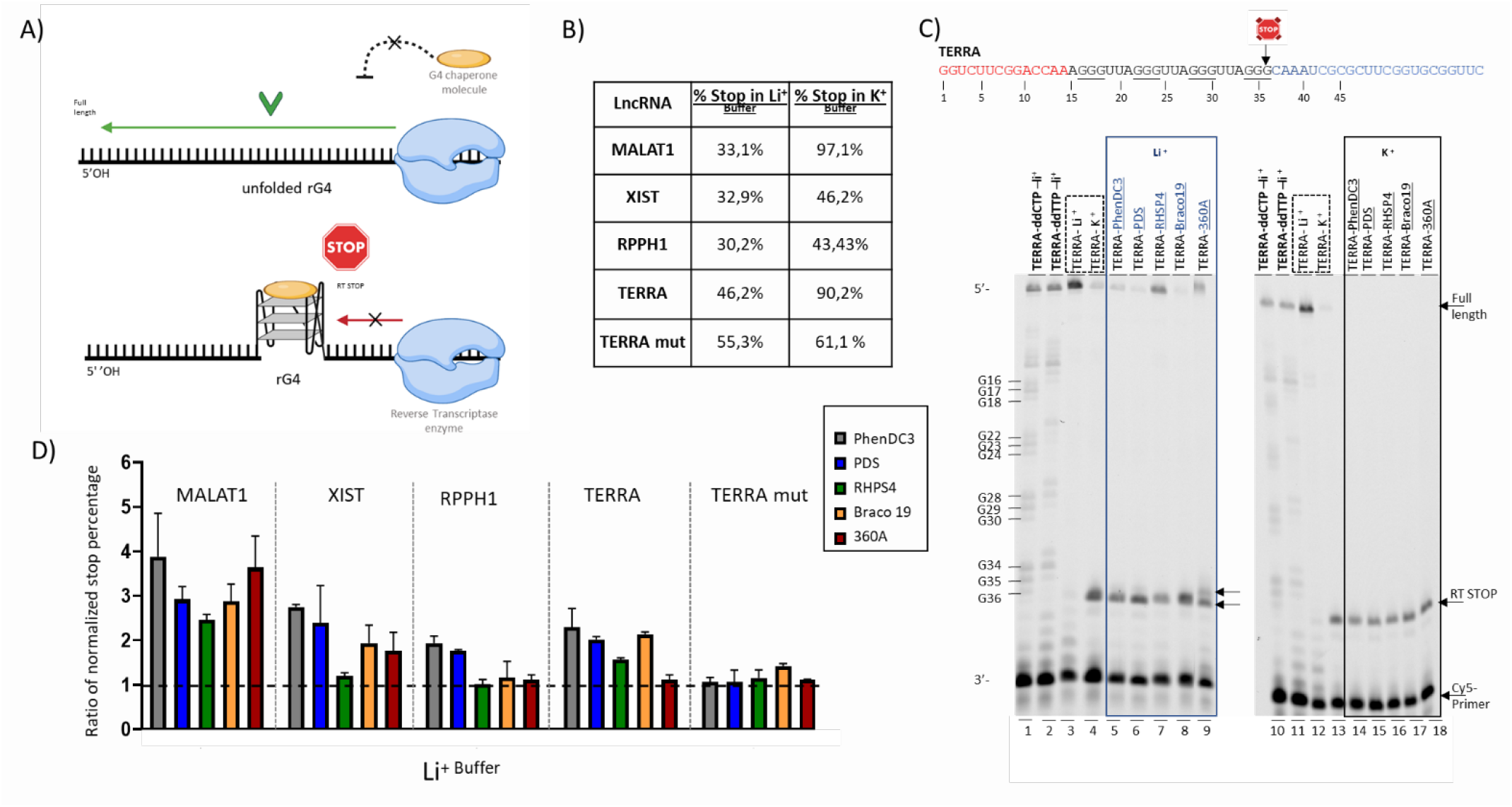
Results from RT stalling assays. A) Schematic representation of the RTS assays used to identify rG4 molecular chaperones. B) Recapitulative results showing the stop percentages obtained by RTS assays in both the Li^+^ and K^+^ conditions without the G4 ligands. C) Example of an RTS assay using TERRA rG4 in the presence of either Li^+^ or K^+^ buffer, without (lines 3 and 4 and lines 12 and 13) or with G4-ligands (*e*.*g*., PhenDC3, PDS, RHPS4, Braco19 and 360A) under either Li^+^ (lines 5,6,7,8 and 9) or K^+^ buffer conditions (lines 14,15,16,17 and 18), respectively. D) Overall results of the RTS assays obtained in the presence of Li^+^ with the four rG4s.

As expected, a higher percentage of stalls was observed in the K^+^ buffer as compared to the Li^+^ buffer, with a more significant gap being observed with both TERRA and MALAT1 (up to 40% and 60%, respectively; see figure 3C and 3D). The reverse transcriptase stalling was observed to be less efficient in the presence of K^+^ buffer for the 2-quartets G4s that form in the RPPH1 and XIST transcripts (13% and 14%, respectively), certainly when compared to the 6% observed with the TERRA mutated sequence, highlighting the ability of these candidates to fold into a G4 in K^+^ buffer without the presence of any ligand despite their lower number of tetrads. The rG4 structures were also confirmed by an enhanced fluorescence signal obtained due to the addition of NMM, a phenomenon that is described as a turn-on parallel G4 probe (Figure S8)^34^. Interestingly, the addition of G4 ligands (10 mol. eq) did not change the stall profiles of MALAT1 and TERRA when the G4s were already folded under these salt conditions (i.e., in K^+^ buffer) in which the G4 formation was optimal (Figure S9). In contrast, in Li^+^ buffer, drastic changes occurred, with some G4-ligands causing the stop of the reverse transcriptase by inducing rG4 formation, which could be considered as them acting like molecular chaperones. Unexpectedly, the stop percentages were only slightly affected by the presence of RHPS4, Braco19, or 360A (i.e., between 1- to 2-fold changes were observed with XIST, RPPH1 and TERRA. In contrast, both PDS and PhenDC3 appeared to be versatile tools, as they induced both classical 3 quartet rG4 structures (TERRA and MALAT1) and non-canonical 2 quartet rG4s (XIST, RPPH1) without any side effects being observed on the control which cannot form an rG4 (TERRA mut). Interestingly, there were no significant increases in the stall with either Braco19 or RHPS4, despite the expectations. Initially Sunny *et al*.^30^ found a negative correlation between the enrichment score change (ΔES) and the corresponding G/C content of the lncRNA transcripts identified by the G4RP-seq. This led them to conclude that sequences with low rG4 formation potential were better structured in the presence of either Braco 19 or RHPS4. Here, a better efficiency of both PDS and PhenDC3 *in vitro* as a molecular chaperone with XIST, MALAT1, RPPH1 and TERRA (Figure 3) was observed. Additionally, the results show that these compounds can thermally stabilize the rG4 in the presence of Li^+^, and that they can accelerate the G4 association process on tetramolecular RNA (Figure 2). Although chaperones can interact with rG4 structures in the transcriptome, and lncRNAs are one possible target, numerous other rG4 structures can be targeted as well. With more than 1.1 million potential structures of rG4, there is a significant diversity among them^48^.

### Exploring the G4-targets of potential molecular chaperones

To investigate the ability of G4 ligands to act as molecular chaperones for a wide range of possible rG4 structures, including both canonical and non-canonical rG4s, a library of synthetic rG4 structures was constructed. This library includes rG4 structures with 2, 3 and 4 tetrads with loops of either 1 or 3 nucleotides between them (Figure 5A). The design of the library was based on a previous study of rG4 thermal stability^49^. The rG4 folding of each synthetic RNA was verified in the presence of either Li^+^ or K^+^ using NMM (See Figure S8). The G4 ligands were then tested using the RTS assays described previously. A heat map of the relative stop percentages of the G4 ligands as compared to the control condition (Li^+^ stop percentage = 1) was generated (Figure 5B).

**Figure 5:**
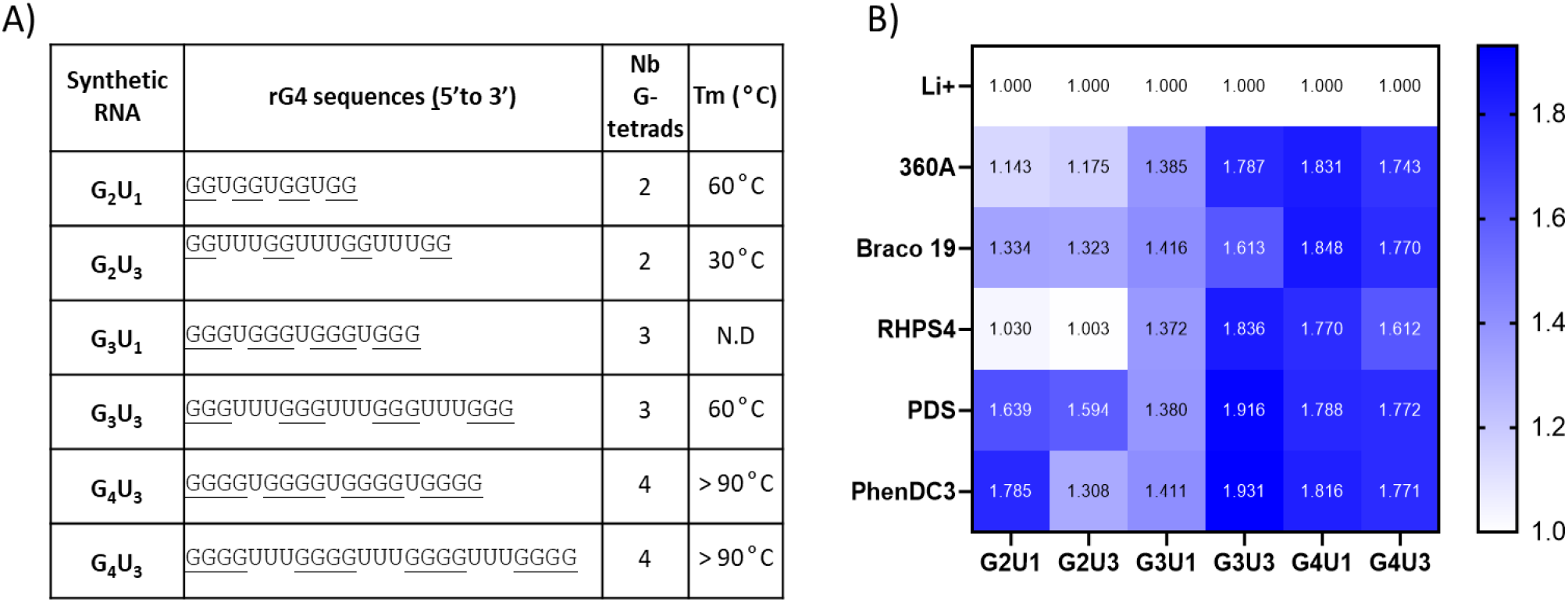
Ligands as molecular chaperone on synthetic rG4. A) A recapitulative table of the synthetic RNA rG4s used in this study, including their numbers of G-tetrads. The stability of this rG4 as determined by Pandley et al^48^ is indicated on the last column of the table. B) Heat maps of the overall quantitative results generates by the RTS assays which were performed in duplicate.

The G4 panels, including Braco 19, RHPS4, PDS, PhenDC3 and 360A, have been confirmed to have inducing activity on very stable structures including 4-quartets (G_4_U_1_ and G_4_U_3_) as well as the 3-quartets (G_3_U_3_). However, RHPS4, BRACO 19 and 360A all exhibit inferior efficiency in inducing the folding of less stable rG4 structures that contain two of the two-quartets (G_2_U_1_ and G_2_U_3_) and one of the three-quartets (G_3_U_1_). The versatility of both PhenDC3 and PDS has been confirmed with both canonical and non-canonical G4 structures, with a mean score of up to 1.38 being obtained. The chaperone rG4 properties of PDS have also been recently raised in a comparative analysis of both the rG4-Seq data and the *in silico* pG4 folding^48^. In fact, it was observed that the number of non-canonical rG4s detected by RTS sequencing significantly increased in all categories (long loop, bulges and 2-quartet rG4s) when the cells were treated with PDS. Notably, a significant increase in 2-quartet rG4 was observed, with over 8 times more RTS sites being detected in PDS-treated cells (5 222) as compared to what was observed in the presence of K^+^ (639)^16^. A study was published in 2012 attempting to assess the impact of PhenDC3 on the transcriptomic landscape, but no in-depth studies on the type of G4 stabilized or induced by this ligand have been reported to date^50^. That said, *in vitro* PhenDC3 has been shown to be highly effective in stabilizing G-tetrads due to its perfectly matched shape, and it has also been reported to induce G4 DNA structure, consistent with our earlier results^32,51^.

It is challenging to conceive how a ligand could induce the formation of a structure on its own without the aid of stabilizing cations. However, a proposal has been suggested that highlights the importance of the PhenDC3 stoichiometry in promoting the formation of dG4 in sequences with low G4Hunter scores. Mass spectrometry experiments reveal that the [G4+2PhenDC3] complex is the most frequently observed species formed from an unstructured strand^52^.

## Conclusions

This study sheds light on the crucial role of some well-known G4-ligands in the formation of G4 structures. Through the evaluation of various panels of G4-ligands, both PDS and PhenDC3 have been identified as specific chaperones that facilitate rG4 formation, and that also stabilize the structures in different RNAs containing from 2 to 4 G-tetrads and derived from lncRNA.

RHPS4, Braco19 and 360A seem to have limited abilities as molecular chaperone on unusual rG4s, while PDS and PhenDC3 are notably more efficient rG4 inducers. One can imagine the weaker G4 ligands more accurately represent G4 formation in cells, while the use of G4 chaperones may lead to the formation of less stable or non-canonical rG4 types that likely cannot form without them. These findings have implications for critically assessing the use of certain ligands as absolute tools for investigating the biology of G4s and for understanding the regulatory mechanisms of RNA species.

Overall, this research highlights the importance of continued investigation into the molecular mechanisms underlying the various roles of G4 ligands. Moreover, these potential roles should be considered when choosing G4 ligands for transcriptome landscape data analysis.

## Acknowledgments

We would like to thank Marc Pirrotta and David Monchaud for their assistance and helpful discussions during the FRET experiments and extend our gratitude to Marc-Antoine Turcotte and Francois Bolduc for their valuable contributions and insightful discussions throughout the entire project.

