## Supplementale Data for "Small molecule chaperones facilitate the folding of long non-coding RNA G"

### Supplementary information

Pauline Lejault, Louis Prudent, Michel-Pierre Terrier, and Jean-Pierre Perreault

1. Oligonucleotides-------------------------------------------------------------Figure S1
2. FRET-melting------------------------------------------------------------------Figure S2
3. EMSA analyses -------------------------------------------------------------Figure S3
4. RTS-quantifications methods----------------------------------------------Figure S4
5. RTS-assay (lncRNA)-------------------------------------------------------Figure S5
6. RTS-assay (Synthetic RNA)--------------------------------------------Figures S6-S7
7. NMM assays ------------------------------------------------------------------Figure S8
8. RTS quantification in presence of $K^+$ -----------------------------------Figure S9

### 1. Oligonucleotides

| Oligonucleotide name (method) | 5'to 3' Sequences | RNA/DNA |
|---|---|---|
| Primer (RTS) | Cy5-[5'GAACCGCACCGAAGCGCG3'] | DNA |
| MALAT1 (RTS) | 5'GAACCGCACCGAAGCGCGATTTGCCCACCCCCTCCTCCCATCCCTTGGTCCGAAGACCTATAGTGAGTCGTATTA3' | DNA |
| TERRA mut (RTS) | 5'GAACCGCACCGAAGCGCGATTTGCCAACCTTCCTTCCTTGGTCCGAAGACCTATAGTGAGTCGTATTA3' | DNA |
| (RTS) | 5'GAACCGCACCGAAGCGCGATTTGCCGCCGGGCCCCTCCTTGGTCCGAAGACCTATAGTGAGTCGTATTA3' | DNA |
| TERRA (RTS) | 5'GAACCGCACCGAAGCGCGATTTGCCCTAACCCTAACCCTAACCCTTGGTCCGAAGACCTATAGTGAGTCGTATTA3' | DNA |
| Ctrl Terra mut (RTS) | 5'GAACCGCACCGAAGCGCGATTTGC**T**CTAAC**T**CTAAC**T**CTAACTCTTGGTCCGAAGACCTATAGTGAGTCGTATTA3' | DNA |
| (G2U1)$_4$ (RTS) | 5'GAACCGCACCGAAGCGCGATTTGCCACCACCACCTTGGTCCGAAGACCTATAGTGAGTCGTATTA3' | DNA |
| (G2U3)$_4$ (RTS) | 5'GAACCGCACCGAAGCGCGATTTGCCAAACCAAACCAAACCTTGGTCCGAAGACCTATAGTGAGTCGTATTA3' | DNA |
| (G3U1)$_4$ (RTS) | 5'GAACCGCACCGAAGCGCGATTTGCCCACCCACCCACCCTTGGTCCGAAGACCTATAGTGAGTCGTATTA3' | DNA |
| (G3U3)$_4$ (RTS) | 5'GAACCGCACCGAAGCGCGATTTGCCCAAACCCAAACCCAAACCCTTGGTCCGAAGACCTATAGTGAGTCGTATTA3' | DNA |
| (G4U1)$_4$ (RTS) | 5'GAACCGCACCGAAGCGCGATTTGCCCCACCCCACCCCACCCCTTGGTCCGAAGACCTATAGTGAGTCGTATTA3' | DNA |
| rA$_{10}$G$_5$A$_{10}$ (EMSA) | 5'AAAAAAAAAAGGGGGA | RNA |
| MALAT1 (FRET) | FAM-[5'GGGAUGGGAGGAGGGGGUGGG3']-TAMRA | RNA |
| TERRA mut (FRET) | FAM-[5'GGAAGGAAGGUUGG3']-TAMRA | RNA |
| RPPH1 (FRET) | FAM-[5'GGAGGGGCCCGGCGG3']-TAMRA | RNA |
| TERRA (FRET) | FAM-[5'GGGUUAGGGUUAGGGUUAGGG3']-TAMRA | RNA |
| F-duplex-T (FRET) | FAM-d[5'TATAGCTATATTTTTTTATAGCTATA3']-TAMRA | DNA |

**Supplementary Table S1. Oligonucleotides used in this study.**
The T7 RNA promoter that was used for *in vitro* transcription is in green. The red nucleotides are those responsible for the 3'-hairpin formation, while those in part are involved in the 5'-hairpin formation. The oligonucleotide tracks which play a role in G-quadruplex folding are underlined.

### 2. FRET melting

#### Supplementary Figure S2: FRET melting curves

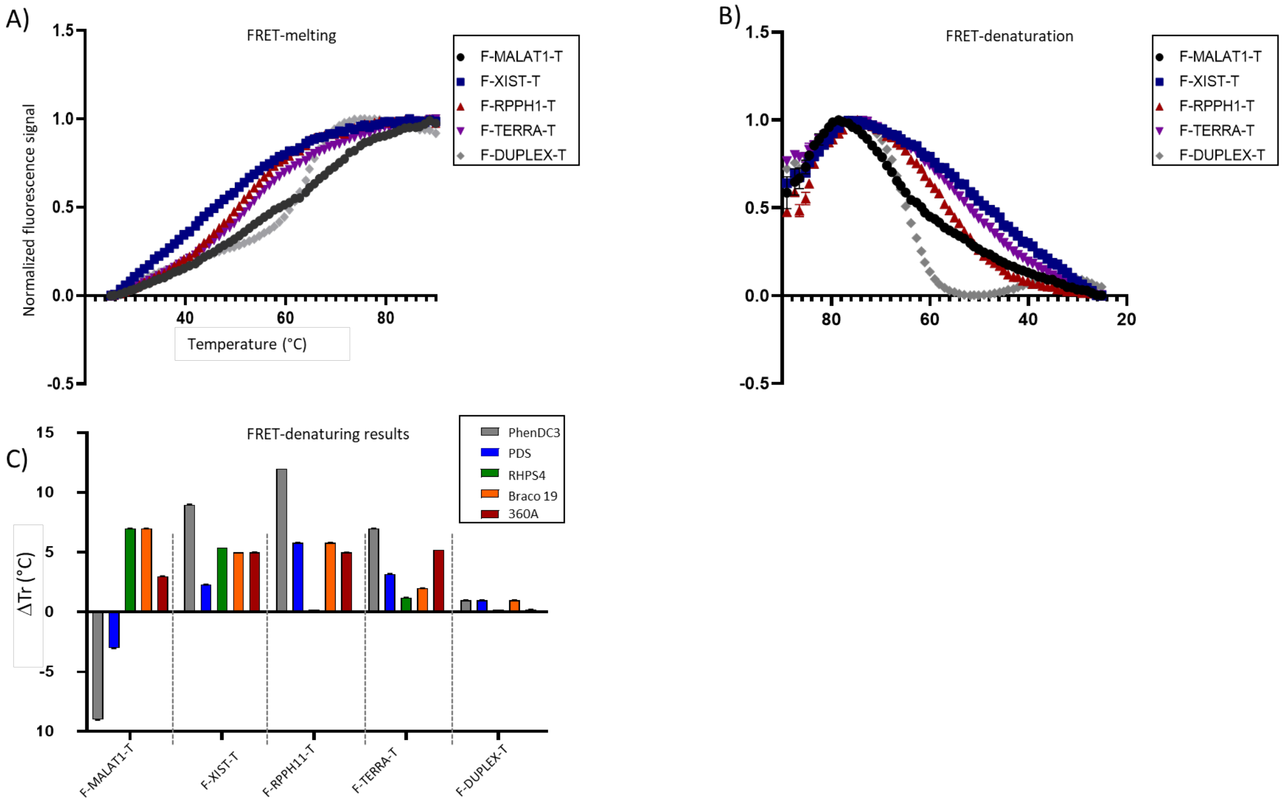


**Supplementary Figure S2. Fret melting/denaturation.** A) FRET melting curves of MALAT1, XIST, RPPH1, TERRA and the DUPLEX control without the presence of any ligands in cacodylate $Li^+$ 1 mM buffer. B) FRET-denaturation curves of MALAT1, XIST, RPPH1, TERRA and the DUPLEX control without the presence of any ligands in cacodylate $Li^+$ 1 mM buffer. C) Overall results of FRET-denaturation with 5 mol. eq of PhenDC3, PDS, RHPS4, Braco 19 and 360A.

### 3. EMSA analyses

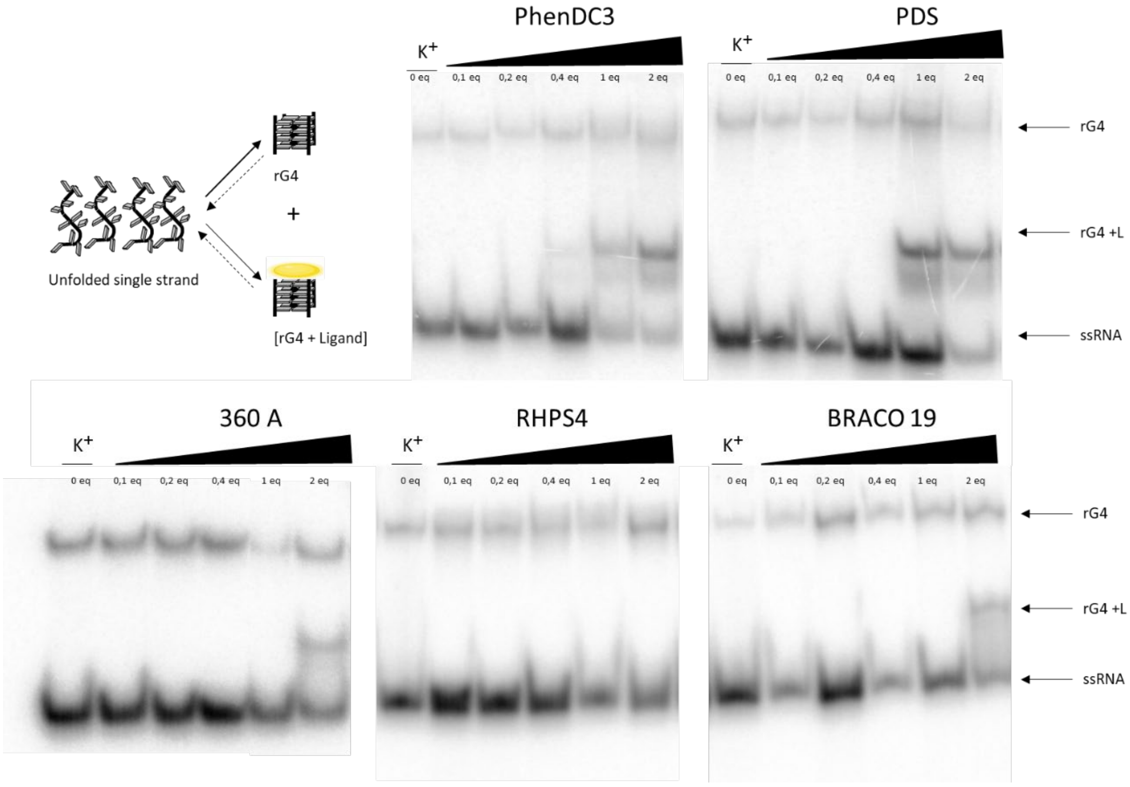


**Supplementary Figure S3: Gel shift assays.** Each experiment was performed with $^{32}P$ labelled-$A_{10}G_4A_{10}$ (20 µM) in the presence of increasing amounts of each ligand (0,1;0,2; 0,4; 1; and, 2 mol. eq). Non-denaturing 20% polyacrylamide gel electrophoresis was performed. The quantification is presented in Figure 2.

### 4. RTS-quantifications methods

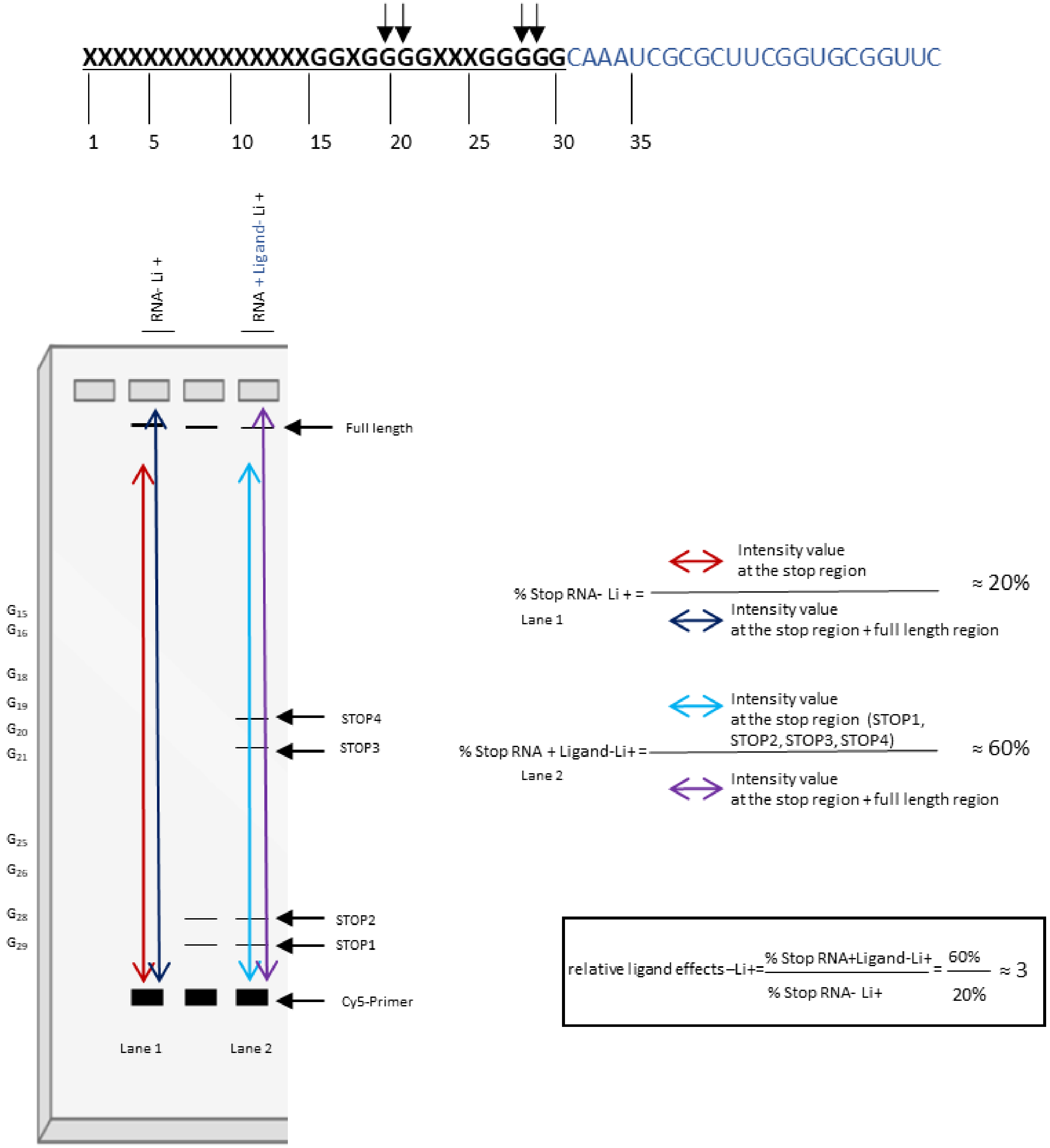


**Supplementary Figure S4: Schematic representation of an RTS-gel and the quantitative methods developed in this study**. The experiment compares the effects of G4-ligands on the ability of rG4 structures to block reverse transcription, measured as %stop RNA. In lane 1, $Li^+$ is used to create a condition where any rG4 structure can block reverse transcription, resulting in a low %stop RNA (20%). In lane 2, the same $Li^+$ condition is used but with the addition of a G4-ligand, which induces rG4 formation and leads to four stops during reverse transcription. This increases the %stop RNA to 60%. To calculate the relative effect of the ligand, divide the %stop RNA-$Li^+$ by the %stop RNA+ligands $Li^+$ to obtain a 3-fold change in the relative ligand effect.

### 5. RTS-assays of rG4 derived from lncRNA

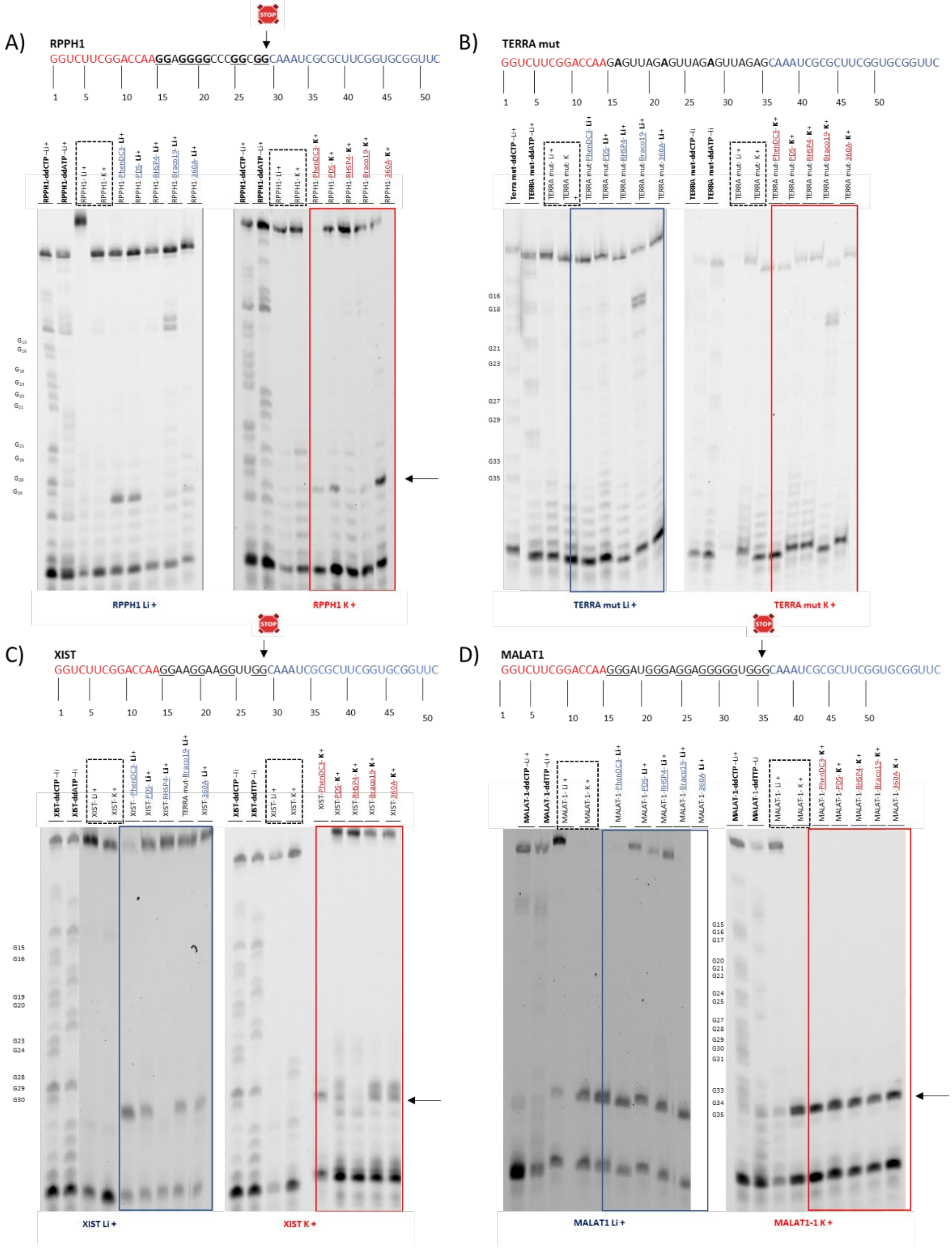


**Supplementary Figure S5: Denaturing polyacrylamide gel electrophoresis of the samples after the RTS-assays.** The experiments were performed with: A) RPPH1, B) Terra-Mut, C) XIST and D) MALAT1. The ddCTP, ddTTP or ddATP ladders indicate the positions of every complementary nucleotide so, every Guanine, Adenine or Thymine, respectively. The RTS assays were performed in the presence of either $Li^+$ (blue box) or $K^+$ ions (red box). The black arrows indicate the major stops involved in the folding the the rG4.

### 6. RTS-assays of synthetic rG4

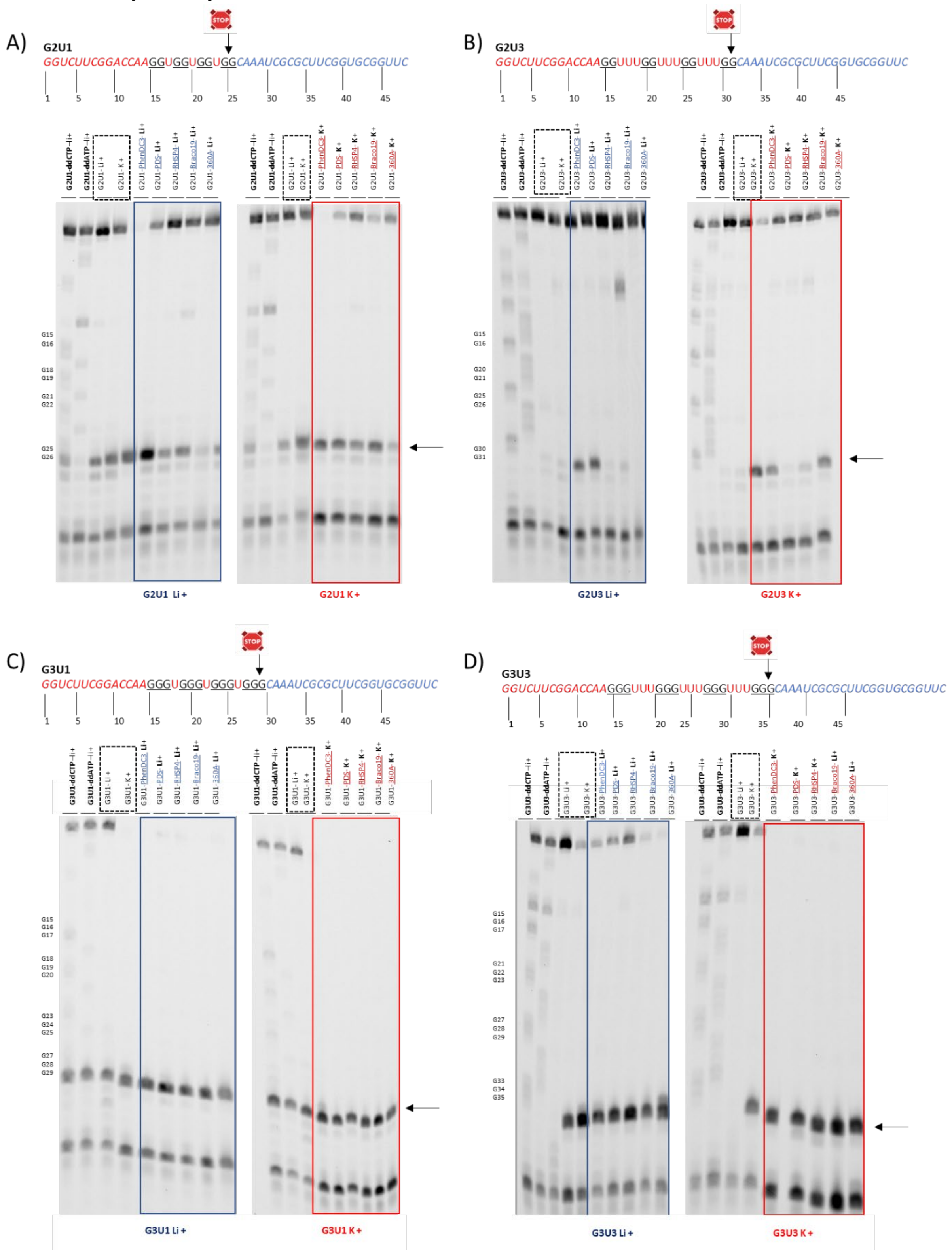


**Supplementary Figure S6: Denaturing polyacrylamide gel electrophoresis of the samples after the RTS-assays.** The experiments were performed with: A) G2U1, B) G2U3, C) G3U1 and D) G3U3. The ddCTP, ddTTP or ddATP ladders indicate the positions of every complementary nucleotide so, every Guanine, Adenine or Thymine, respectively. The RTS-assays were performed in the presence of either $Li^+$ (blue box) or $K^+$ ions (Red box). The black arrows indicates the major stops involved in the folding the the rG4.

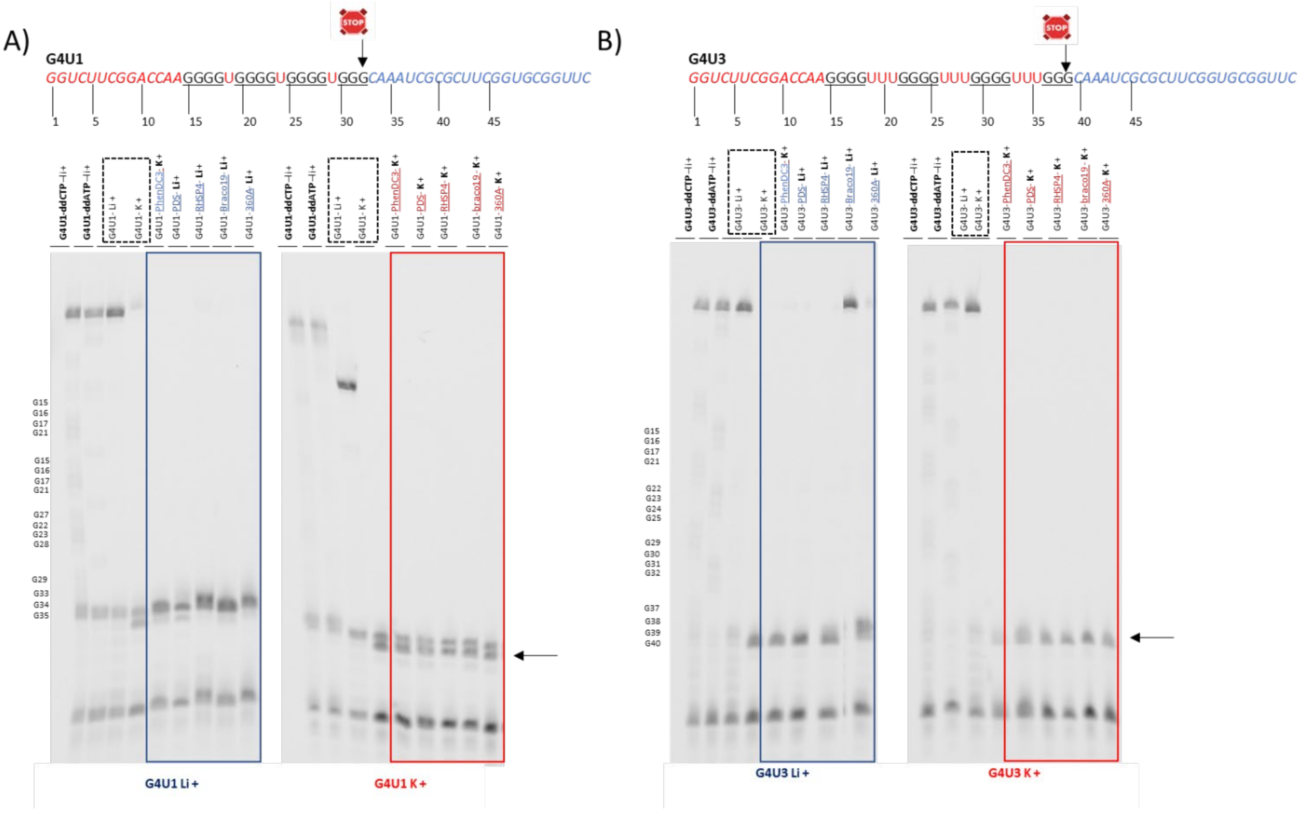

**Supplementary Figure S7: Denaturing polyacrylamide gel electrophoresis of the samples after the RTS-assays.** The experiments were performed with: A) G4U1 and B) G4U3. The ddCTP, ddTTP or ddATP ladders indicate the positions of every complementary nucleotide so, every Guanine, Adenine or Thymine, respectively. The RTS assays were performed in the presence of either $Li^{+}$ (blue box) or $K^{+}$ ions (Red box). The black arrows indicate the major stops involved in the folding the G-quadruplex.

### 7. NMM assays

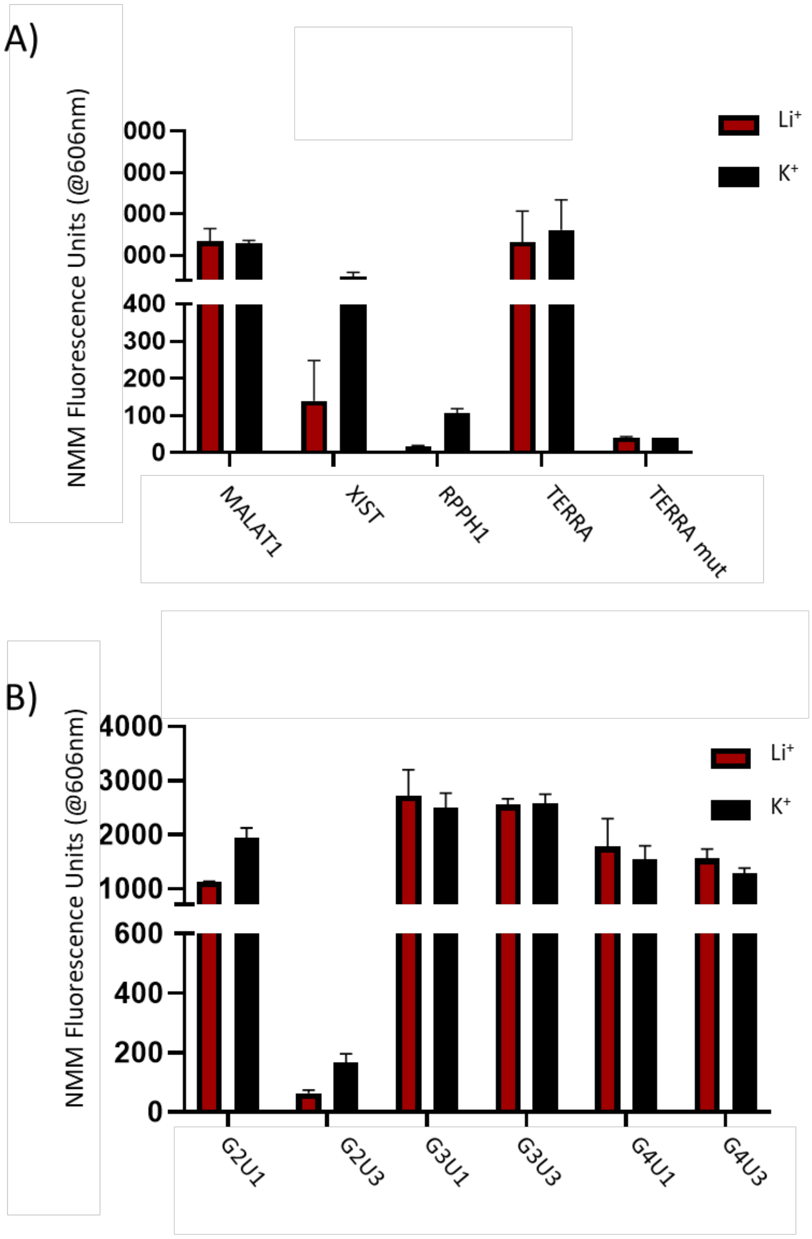


**Supplementary Figure S8: N-methyl mesoporphyrin (NMM) fluorescence assays.** The histograms of the fluorescence emissions (at 606 nm) tested with the *in vitro* transcribed RNAs used in the RTS assays are shown with: A) MALAT1, XIST, RPPH1 and TERRA rG4; and, B) G2U1, G2U3, G3U1, G3U3, G4U1 and G4U3 synthetic rG4. The different salt conditions are indicated as follows: $Li^+$ in red and $K^+$ in black. Each bar represents the mean of 2 independent experiments, and the error bars represent the standard deviations.

### 8. RTS quantification in the presence of K+

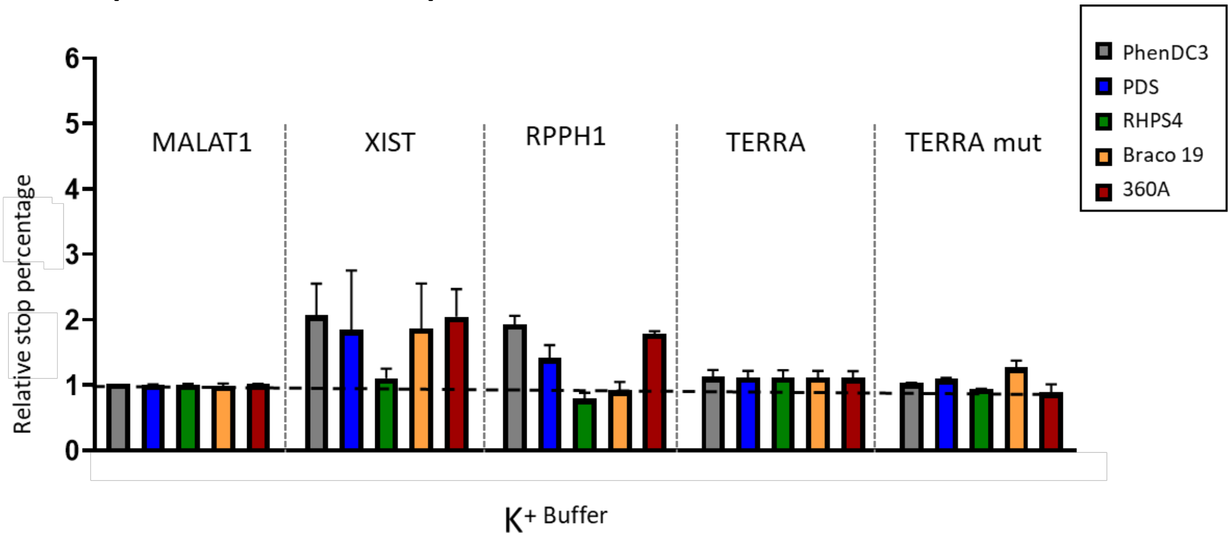


**Supplementary Figure S9: Overall results of the RTS assays in the presence of $K^+$ with the four rG4 derived from the lncRNAs (MALAT1, XIST, RPPH1, TERRA, TERRA mut)** with 10 mol. eq of PhenDC3, PDS, RHPS4, Braco 19 and 360A.

.
